# Lesion site and therapy time predict responses to a therapy for anomia after stroke: a prognostic model development study

**DOI:** 10.1101/2021.02.05.429894

**Authors:** Thomas M.H. Hope, Davide Nardo, Rachel Holland, Sasha Ondobaka, Haya Akkad, Cathy J. Price, Alexander P. Leff, Jenny Crinion

**Affiliations:** Institute of Cognitive Neuroscience, University College London, UK; Wellcome Centre for Human Neuroimaging, University College London, UK; MRC Cognition and Brain Sciences Unit, Cambridge University, UK; Division of Language and Communication Sciences, City University London, UK; UCL Queen Square Institute of Neurology, London, UK

## Abstract

**BACKGROUND:** Stroke is a leading cause of disability, and language impairments (aphasia) after stroke are both common and particularly feared. Most stroke survivors with aphasia exhibit anomia (difficulties with naming common objects), but while many therapeutic interventions for anomia have been proposed, treatment effects are typically much larger in some patients than others. Here, we asked whether that variation might be more systematic, and even predictable, than previously thought.

**METHODS:** 18 patients, each at least 6 months after left hemisphere stroke, engaged in a computerised treatment for their anomia over a 6 week period. Using only: (a) the patients’ initial accuracy when naming (to-be) trained items; (b) the hours of therapy that they devoted to the therapy; and (c) whole-brain lesion location data, derived from structural MRI; we developed Partial Least Squares regression models to predict the patients’ improvements on treated items, and tested them in cross-validation.

**RESULTS:** Somewhat surprisingly, the best model included only lesion location data and the hours of therapy undertaken. In cross-validation, this model significantly out-performed the null model, in which the prediction for each patient was simply the mean treatment effect of the group. This model also made promisingly accurate predictions in absolute terms: the correlation between empirical and predicted treatment response was 0.62 (95%CI: 0.27, 0.95).

**DISCUSSION:** Our results indicate that individuals’ variation in response to anomia treatment are, at least somewhat, systematic and predictable, from the interaction between where and how much lesion damage they have suffered, and the time they devoted to the therapy.

## 1. Introduction

Stroke is a leading cause of disability [1], and language impairments (aphasia) after stroke are both common [2] and particularly feared [3]. Most stroke survivors with aphasia exhibit anomia, a difficulty finding words when naming common objects [4], but while many therapeutic interventions for anomia have been proposed, treatment effects typically vary, substantially, from patient to patient [5]. Inter-individual variation in treatment responses is ubiquitous in medicine (e.g. in psychiatry and pharmacology, respectively [6, 7]), but emerging evidence suggests that variation in responses to therapy for aphasia after stroke might be more systematic than previously thought [5, 8, 9]. To the extent that this is true, the implication is that we can use pre-treatment (e.g. behavioural and / or brain imaging) data to explain and even predict patients’ likely treatment responses.

For example, we recently showed that a model derived from: (a) pre-treatment scores on standardised cognitive and language tasks; and (b) lesion location data, derived from pre-treatment structural MRI, could be used to predict 23 aphasic stroke patients’ responses to a treatment for acquired reading impairments (central alexia) [8]. Like the treatment considered here, for anomia, this earlier treatment for alexia was a computerised application designed to engage participants in massed practice of trained items at home, over a period of weeks. Using stepwise forward feature selection, we selected specific predictors from the pre-treatment data for entry into a multiple linear regression model, which explained over 90% of the variance in patients’ empirical treatment responses. This result is biased by over-fitting, because all of the patients’ data were used to select features, but even when the feature selection was nested within each fold of a leave-one-out cross-validation process (i.e. removing the bias), the resulting predictions were significantly correlated with patients’ empirical treatment responses (r = 0.48, p < 0.05) [8].

In what follows, we add to the evidence that responses to aphasia treatment might be predictable from patients’ pre-treatment data. Here, we focus on a computerised treatment for naming difficulties (anomia) that is the most common language impairment after stroke [4]. This treatment’s effectiveness at the group level has already been verified [10]; here, we attempt to explain and predict the same treatment effects at the individual level. Our main hypothesis was that the individual patients’ responses to the treatment are systematic and predictable given where and how much lesion damage they had suffered. We tested this by comparing predictions made by models derived from those data (alone or in combination with their pre-treatment anomia severity, demographic data and the hours that they devoted to the therapy), to the predictions made by a ‘null’ model, which simply predicts the mean treatment response of its training sample. If (any of) the former are significantly more accurate than the latter, the implication is that individual variation in responses to this treatment is systematic and predictable, at least to some extent.

## 2. Methods

The current analysis employs pre-treatment: (a) demographic data (age at stroke onset, time post-stroke at assessment, and sex); (b) patients’ initial impairment severity (i.e. their accuracies when naming to-be-treated items, pre-treatment; (c) the hours of therapy actually undertaken; and (d) structural MRI, which we use to extract lesion location data. We use these data to predict and explain patients’ responses to therapy, measured as the absolute change in naming accuracy, from pre-to post-treatment, for ‘trained’ items (i.e. items practiced during the therapy).

The therapy was designed to engage participants in massed practice of object naming, over a 6 week period at home. A variety of phonemic cues (e.g. an audio recording of the object’s name, or of the name’s first phoneme) were presented concurrently with the picture to be named during treatment to encourage error-reducing learning. The approach was both effective and specific to spoken word production, significantly improving patients’ overall object naming accuracy and reaction time immediately post-treatment (unstandardized effect size: 29% and 17%, respectively; Cohen’s *d*: 3.45 and 1.83). Longer term gains in naming were maintained three months later, though in this study we focus only on the immediate gains made for items trained during the therapy.

### 2.1 Participants

The study participants were 18 right-handed native English-speakers, with normal hearing, no history of psychiatric disease and no prior history of neurological disorder before suffering a left-hemisphere stroke, causing language impairment (aphasia). Participants were recruited either from an aphasia clinic, run by JC, or via the Predicting Language Outcomes After Stroke (PLORAS) study, run by CJP, between 2009 and 2012. The study size was arrived at via a power calculation based on the expected effect size of the treatment considered.

Participants were only included if they had: (i) naming difficulties (anomia), as assessed via the Boston Naming Test (cut-off <56); (ii) relatively preserved single-word comprehension as assessed via the Comprehensive Aphasia Test (CAT) [11]; (iii) good mono-syllabic word repetition as assessed via the Psycholinguistic Assessments of Language Processing in Aphasia [12]; (iv) no speech apraxia as determined by the Apraxia Battery for Adults [13]; and, (v) at least partially spared left inferior frontal cortex (thought to support speech re-learning [10]). All gave written informed consent to take part in the study, which was approved by the Central London Research Ethics Committee and conducted in accordance with the ethical principles stated by the Declaration of Helsinki. A table of the participants’ key characteristics, reproduced from [10], is included in supplementary material.

### 2.2 Stimuli and procedure

The procedure for the treatment study [10] involved behavioural assessments and neuroimaging data acquisition both pre- and post-treatment. Here, we use pre-treatment data only, to predict treatment response, calculated as the change in the number of trained items that patients could name correctly.

Stimuli were drawn from a pool consisting of 299 black and white line drawings of objects adapted from the International Picture-Naming Project (https://crl.ucsd.edu/experiments/ipnp/). All object names were monosyllabic, with a consonant-vowel-consonant structure and high name agreement (e.g. ‘car’). The treatment employed 150 of the 299 stimuli: i.e. for each patient, there were 150 treated items and 149 untreated items. 54/150 to-be-trained items and 53/149 un-trained items were kept common across all patients (for use in an FMRI experiment [10], which we do not consider here). The remaining items (96/150 to be trained; 95 /150 to be untrained) were determined for each patient on the basis of their individual pre-treatment naming performance (accuracy) on the 299 items, to match each patient’s pre-treatment performance on treated and untreated lists, respectively.

After baseline assessment and pre-treatment structural MRI, patients were given a laptop and asked to complete a minimum of two hours of naming practice 5 days a week, over a six-week period. The pictures and auditory cues were presented using the ‘StepByStep’ aphasia treatment software (http://www.aphasia-software.com). The naming practice was designed to be completed in an error-reducing manner [14]. For example, in naming a picture of a car the patient was asked to name it three times: (i) after a whole word auditory cue /ka:r/; (ii) after an initial phonemic cue /ka/; (iii) after a whole word cue again. Only then would the patient proceed to the next item to be named. Patients completed on average a total of 73 (± 25) hours of naming practice: i.e. within one standard deviation of the mean therapy dose found, in a meta-analysis of aphasia treatment studies [15], to improve patients’ communicative ability. After the six-week period, patients were assessed again exactly as at baseline. Naming accuracy was scored according to the standardized Comprehensive Aphasia Test guidelines [11]. Our analyses here are separately focused on absolute change in naming accuracy (from pre-to post-treatment) on the 150 treated items.

### 2.3 Imaging acquisition and analysis

The same scanner and hardware were used for the acquisition of all images. Whole-brain imaging was performed on a 3 T Siemens TIM-Trio system (Siemens) at the Wellcome Centre for Human Neuroimaging. Lesion images were derived from structural MRI using the Automatic Lesion Identification toolbox [16], and then double-checked for accuracy by a researcher experienced in manual lesion-tracking (DN),, working on individual axial slices.

Lesion data were then encoded as lesion load in a series of 398 anatomically defined regions of interest, derived from four publicly available atlases (two focused on grey matter and two focused on white matter) [17-20]. Where regions were represented in probabilistic format, they were re-encoded as binary images at a 50% threshold. For each region, lesion load was calculated as the number of (2mm^3^) voxels shared by the lesion and the region, divided by the total number of voxels in that region. Notably, there was significant overlap between these regions, across atlases. Rather than deciding *a priori* what the best or most useful atlas might be, our goal was simply to reduce the dimensionality of the lesion data in a manner that retained an explicit link with familiar brain regions and / or tracts.

### 2.4 Modelling Methods

Our key aim here was to assess whether individual patients’ treatment responses could be predicted from pre-treatment data alone. Here, we define ‘treatment responses’ as the absolute change in patients’ naming accuracies from pre-to post-treatment.

Treatment studies in this domain are resource intensive and typically involve massed practice, so take time to complete. Like most others in the field, our sample is therefore smaller (n=18) than is usually desirable when building predictive models, increasing the risk of over-fitting. That risk is further increased because we have so much pre-treatment data to consider, including behavioural data, and lesion data derived from structural MRI.

One way to manage this risk is via feature selection, as we employed in similar, previous work [8]. But though successful, that work still revealed significant over-fitting, because our in-sample results (using the whole dataset to select features) were so much stronger than our out-of-sample results (i.e. nesting feature selection in cross-validation): R^2^ (predicted response, empirical response) = 0.94 (in-sample); 0.23 (out-of-sample). Accordingly, we took a simpler approach in this work by using dimensionality reduction, rather than feature selection, to manage the high dimensionality of the pre-treatment (behavioural and brain imaging) predictors.

#### 2.4.1 Predictive models

We used Partial Least Squares (PLS) regression, as implemented in Matlab 2019a, to develop our models, using either: (a) demographic variables, including age at stroke onset, sex and time post-stroke; (b) pre-treatment naming accuracy (i.e. measuring the initial severity of each patient’s anomia); and / or (c) lesion data, derived from pre-treatment structural MRI. We additionally considered one further variable, both singly and in combination with the other data: the hours of therapy actually completed by each patient. There were no missing data for any patient for any of these variables. All predictor variables were standardised (z-scored) prior to entry into models.

PLS regression is appropriate, here, because it employs dimensionality reduction analogous to, but more efficient than, that implemented by Principal Components Analysis (PCA): i.e. where PCA identifies latent variables which explain maximal variance in the predictors, PLS regression identifies variables that explain maximal variance in the response variable(s). PLS regression thus allows us to build potentially effective models that are (at least somewhat) robust to irrelevant predictors, rather than excluding those predictors explicitly.

The behavioural model employed 28 predictors: i.e. scores on our battery of pre-treatment language and cognitive assessments (as described in detail in [10]). The lesion data were encoded as described previously, into 398 lesion load variables: however, all patients had left-hemisphere lesions, and in fact all patients had zero lesion load in 220/398 regions. These were removed from the analysis (leaving 178 variables), but their removal had no substantive effect on the results. We trained models employing predictors derived from each data type separately, and all higher order combinations of data types.

#### 2.4.2 Model assessment and model comparison

Predictive performance was assessed with cross-validation. We report results using 1,000 times 10-fold cross-validation here, but analyses employing different types of cross-validation were substantially similar. Absolute measures of predictive performance are suspect in small samples, so we assessed our models in relative terms, by comparing them to an empty, or baseline (i.e. null) model, which simply predicts the average treatment response for all patients in the group. This model reflects our null hypothesis, that treatment responses are *not predictable at the individual level*, leaving group-level averages as the only recourse. When our empirical models outperform the empty model, we reject the null hypothesis, concluding that individual treatment responses are predictable, at least to some extent. We compare models by recording the Root Mean Squared Error (MSE) for predictions made in each of 1,000 repetitions of a 10-fold cross-validation. These folds are kept identical across models, so the MSE values can be compared pair-wise.

Traditional paired tests are not appropriate on their own here because different partitions of the data will create overlapping training datasets, which are therefore not independent of each other. Accordingly, while we use the traditional, paired, non-parametric Wilcoxon signed rank test to compare MSEs across models, we further threshold those statistics with paired permutation test. The test construes the two vectors to be compared as having labels, reflecting the models used to generate them. The null hypothesis is that those labels are arbitrary, because the models’ performance do not differ except by chance. We therefore create a null distribution of paired test statistics by randomly permuting those labels *within each pair*, and repeating the original paired (signed rank) test. If the original statistic is extreme relative to the null distribution, we conclude that the performance difference between the models is significant (p < 0.05) after a correction for FamilyWise Error (FWE).

#### 2.4.3 Model interpretation

PLS regression models can be interpreted by examining the weights of each of their components on each of the original variables. However, this approach can be challenging when there are multiple components to consider, and because the sign of each component is arbitrary: i.e. positive weights on a given component do not necessarily imply a positive relationship between the highly weighted independent variables and the dependent variable(s). We circumvent these issues with ‘data perturbation’.

The data perturbation procedure involves permuting random subsamples of the empirical independent variables and recording the effect of the perturbation on the model’s predictions. The PLS model beta weights themselves are fixed based on the original empirical data: our goal is not to fit further models, but rather to better understand the relationships that have already been encoded. We do this by: (a) permuting random subsets of the independent variables; (b) observing the effect on the models’ predictions; and (c) relating perturbed variable values to the resulting predictions with mass univariate correlation analyses. The resultant correlation coefficients approximate the influence that each variable has on the model’s predictions. We ran 1,000 iterations of the process per model, yielding a total sample size of 18 (patients) * 1,000 (iterations), including both perturbed independent variables and the resultant, predicted dependent variable (treatment response). Repeated analyses with this number of iterations yielded very consistent coefficients for all of the models we report across ten repetitions of 1,000 iterations of this process, all pairwise correlations between derived weights on behavioural and lesion variables were >0.99.

## 3. Results

### 3.1 Predictive performance

Table 1 reports predictive performances (median and inter-quartile ranges of Mean Square Errors, or MSEs) of models driven by all combinations of the data we considered. All but one of the models that out-performed the null model, with lower MSEs, included lesion data. The exception was a model including hours of therapy only, with a median MSE of 300 (IQR = 16): i.e. a very small difference relative to the null model, albeit a significant one (FWE adjusted p < 0.05). The best combination was hours of therapy plus lesion data (MSE median / inter-quartile range = 182 / 21), and indeed this was the only combination which improved upon lesion data alone: see Table 1. The mean predictions of that best model, across the 1,000 repetitions, were strongly and significantly correlated with empirical treatment responses (r = 0.62, p = 0.006, 95% CI = 0.27, 0.95): see Figure 1.

**Table 1:** Data configurations and predictive performance, as assessed across the same 1,000 10-fold cross-validation runs. MSE = Mean Squared Errors of the model predictions; IQR = Inter-Quartile Range of the model predictions. These quantities are employed in preference to mean and standard deviation because MSEs typically have a Poisson distribution rather than a normal distribution. Lower MSEs imply more accurate predictions. The best model configuration is underlined (Hrs + Lesions): the most accurate predictions are derived from these data.

| Data types | Median / IQR MSE |
| --- | --- |
| Null | 303/16 |
| Hrs (Therapy) | 300/16 |
| Initial (severity) | 396/31 |
| Demographics | 321/24 |
| Lesions | 205/30 |
| Initial + Hrs | 364/30 |
| Demographics + Hrs | 316/26 |
| **<u>Lesions + Hrs</u>** | **<u>182/21</u>** |
| Initial + Demographics | 364/30 |
| Initial + Lesions | 253/29 |
| Lesions + Demographics | 267/21 |
| Hrs + Initial + Demographics | 355/30 |
| Hrs + Initial + Lesions | 220/32 |
| Hrs + Demographics + Lesions | 253/21 |
| Initial + Demographics + Lesions | 274/23 |
| Hrs + Initial + Demographics + Lesions | 261/23 |

**Figure 1:**
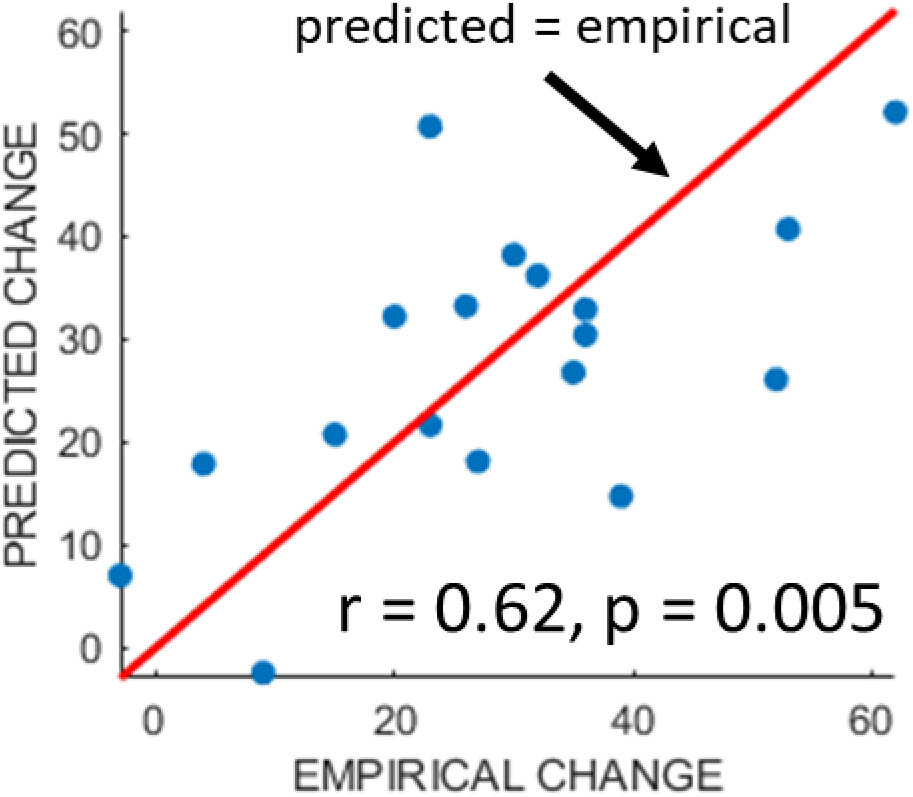
Predicted responses versus empirical responses, for the best model identified in Table 1 (lesion load variables + hours of therapy undertaken).

### 3.2 Interpreting the best model

Variable weights for the best models predicting change on treated items, using a combination of the hours of therapy undertaken and lesion location data, were calculated via data perturbation, as described in the Methods.

First, as expected, the best model predicted better improvement when patients devoted more hours to practice (r = 0.33). Regional weights for the lesion data in this model (i.e. taking therapy hours into account) are displayed in Figure 2, with the most negative weights (predicting lesser treatment benefit with more damage) in and around the left inferior frontal gyrus, and positive weights (predicting greater treatment benefit with more damage) in the middle, superior and anterior temporal lobe regions. Where voxels appear in two overlapping regions with different weights (e.g. we had one region covering the whole of the hippocampus and others covering only its cornu ammonis and dentate gyrus subfields), the most extreme of those two weights is displayed.

**Figure 2:**
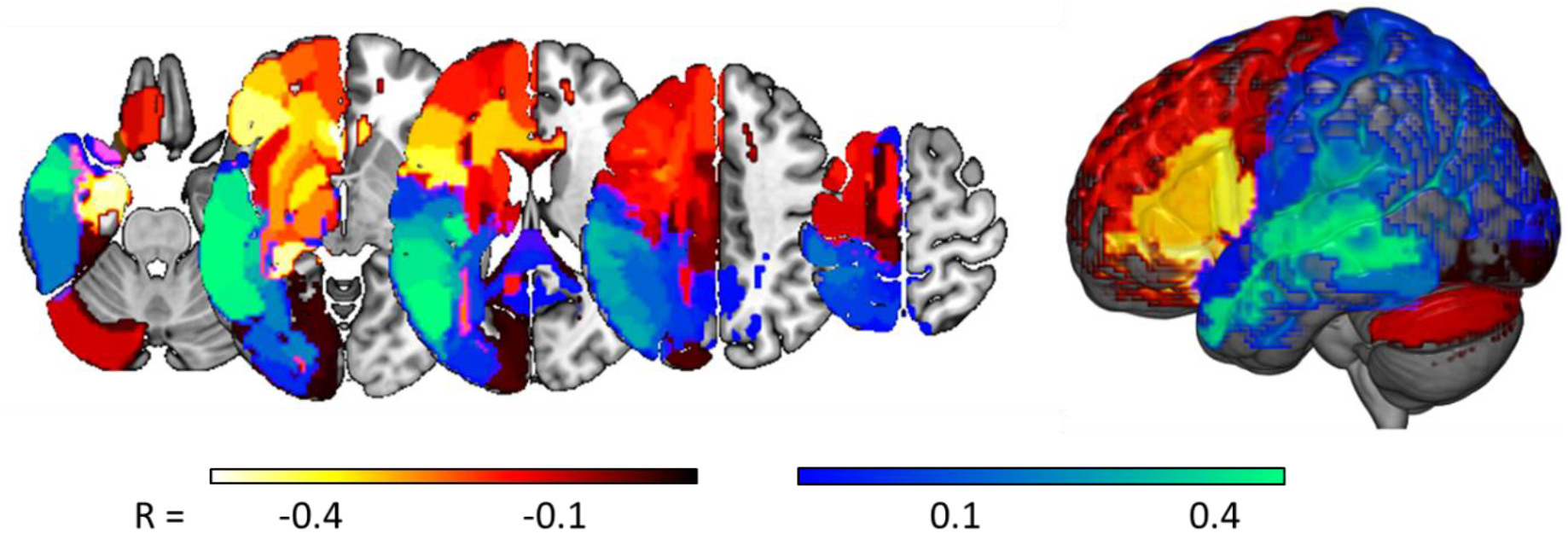
Relating lesion locations to predicted treatment responses. Correlation coefficients, derived via data perturbation, relating the lesion load in each of 177 regions, to treatment responses predicted by our best model (appending lesion data to the hours of therapy actually undertaken).

Notably, weights in many brain areas, including the auditory cortex and the superior, middle and anterior temporal lobes, are positive. The potentially curious implication here, is that more damage predicts larger treatment responses. Instead, we suggest that these positive regions are driven by the contingent distribution of the patients’ lesions: more damage in those positively weighted regions implies less damage in the negatively weighted regions (where the latter make the more intuitive association between more damage and smaller treatment responses). As an illustration of this relationship, we considered area TE11 of the primary auditory cortex, where the strongest, positive weight was observed (0.47). Pairwise correlations, between lesion loads in this region and lesion loads in each of the other (177) regions under consideration, were very strongly correlated with the weights displayed in Figure 2, which were assigned to those regions by our best PLS regression model (r = 0.90): see Figure 3.

**Figure 3:**
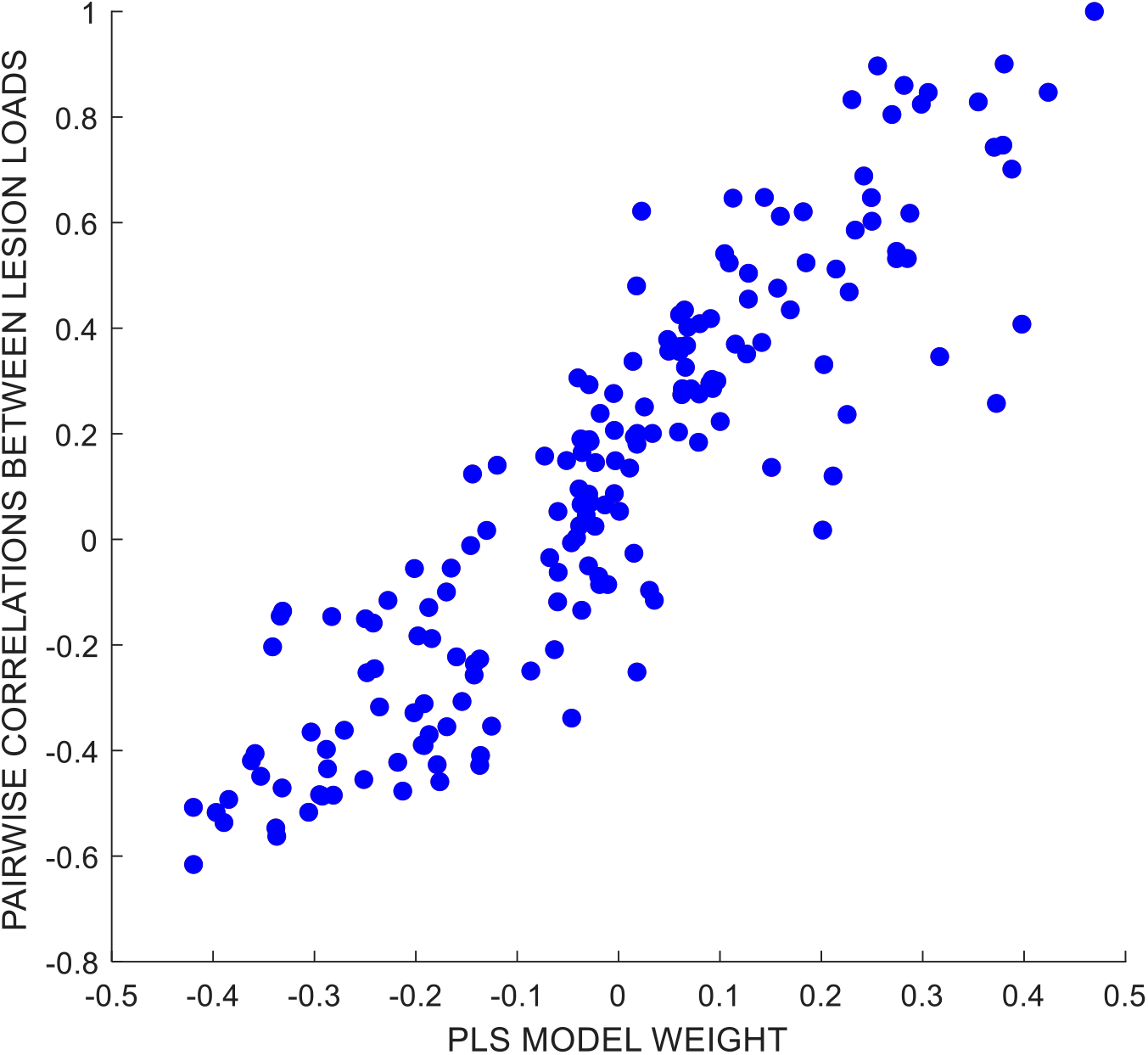
Scatter plot relating: (i) coefficients of the pairwise correlations between lesion load values in primary auditory cortex area TE11, and lesion loads in all of the 177 brain regions that we considered (y-axis); to (ii) the weights assigned to each of those same brain regions by our best PLS regression model, as derived via data perturbation (described in the Methods). The strong correlation between these two quantities implies that lesser lesion load in primary auditory cortex area TE11 serves as a proxy for greater lesion load in areas where that extra damage most strongly predicts poorer treatment responses.

## 4. Discussion

Recent results suggest that individual stroke patients’ responses to aphasia treatment are to some extent systematic and predictable [8]. Our results add to this evidence, showing that responses to a behavioural treatment for anomia are at least somewhat predictable at the individual level. We assessed models derived from demographic variables, and from pre-treatment behavioural and lesion data. Models were derived via PLS regression, drawing on efficient predictor dimensionality reduction and thus obviating the need for either algorithmic or *a priori* feature selection. These models provide a sound way to establish at an individual level whether pre-treatment data include signals that might be used to predict treatment responses.

Many of the models we tested made significantly better predictions than those of a baseline model, in which each patient’s prediction was simply the mean response of the group (see Table 1). However, the best model employed lesion location data, derived from MRI, plus the hours of therapy undertaken by each patient. As hours of therapy alone has very little predictive power, the results suggest that the benefit of increased therapy depends on lesion location. This may explain why detecting therapy dose effects has been so challenging [21]. Notably, we could not predict training effort, as indexed by hours of therapy undertaken at the individual level, from any of the other data considered here.

Our best prognostic model, including lesion data and hours of therapy, is broadly sensible. The negative weights assigned to the left inferior frontal gyrus, the hippocampus and the cerebellum (more damage = less improvement) are consistent with prior work emphasising the importance of the preservation of these regions in the response to aphasia therapy (e.g. [22-24]). And the positive weights may best be explained as emphasising those regions where more extensive damage predicts better preservation of the regions that appear to support better responses to treatment (see Figure 3).

Notably, we did not employ any feature selection in this work: i.e. we did not attempt to select the subset of lesion location variables that might best explain the patients’ treatment responses. This is a limitation of the current work, made necessary because feature selection encourages over-fitting in small samples [8]: the only general way to circumvent this issue is via external validation: testing the best model from this study in a second, completely independent sample. But this is no simple endeavour, because the time and effort required to run these studies is substantial, and we do not yet know how similar such a study would need to be to that reported here. Does the treatment have to stay exactly the same? How much can the inclusion criteria vary? Work to address these questions, by measuring how prognostic models generalise across independent samples (e.g. as in [25]) and different therapy studies, is ongoing.

Perhaps surprisingly, our models did not benefit from the addition, either of the initial severity or the demographic data that we considered – suggesting that this treatment’s efficacy did not depend on the patients’ ages, sex, time post-stroke or pre-treatment impairment severity (once lesion location had been taken into account). Whether these null results generalise in larger samples, is a question for future work. But our results do suggest that pre-treatment structural neuroimaging (lesion data), in combination with treatment dose, can be used to predict individual patients’ therapeutic anomia intervention response. This is consistent with prior results, suggesting that the individual responses to treatment for aphasic stroke might interact with where and how much lesion damage individual patients have suffered [8]. We hope that these results will encourage further attempts to explain and predict inter-individual differences in treatment responses, with pre-treatment data, opening the way for a more positive and personalised treatment approach for aphasia.

## ACKNOWLEDGMENTS

This study was supported by Wellcome (203147/Z/16/Z; 205103/Z/16/Z; 106161/Z/14/Z), MRC (G0701888), the Stroke Association (TSA PDF 2017/02) and NIHR (RP-2015-06-012). The funders had no participation in the design and results of this study.

## DATA

The data described in this study is available to accredited researchers from JC, on request.

---

**Supplementary Table S1:**
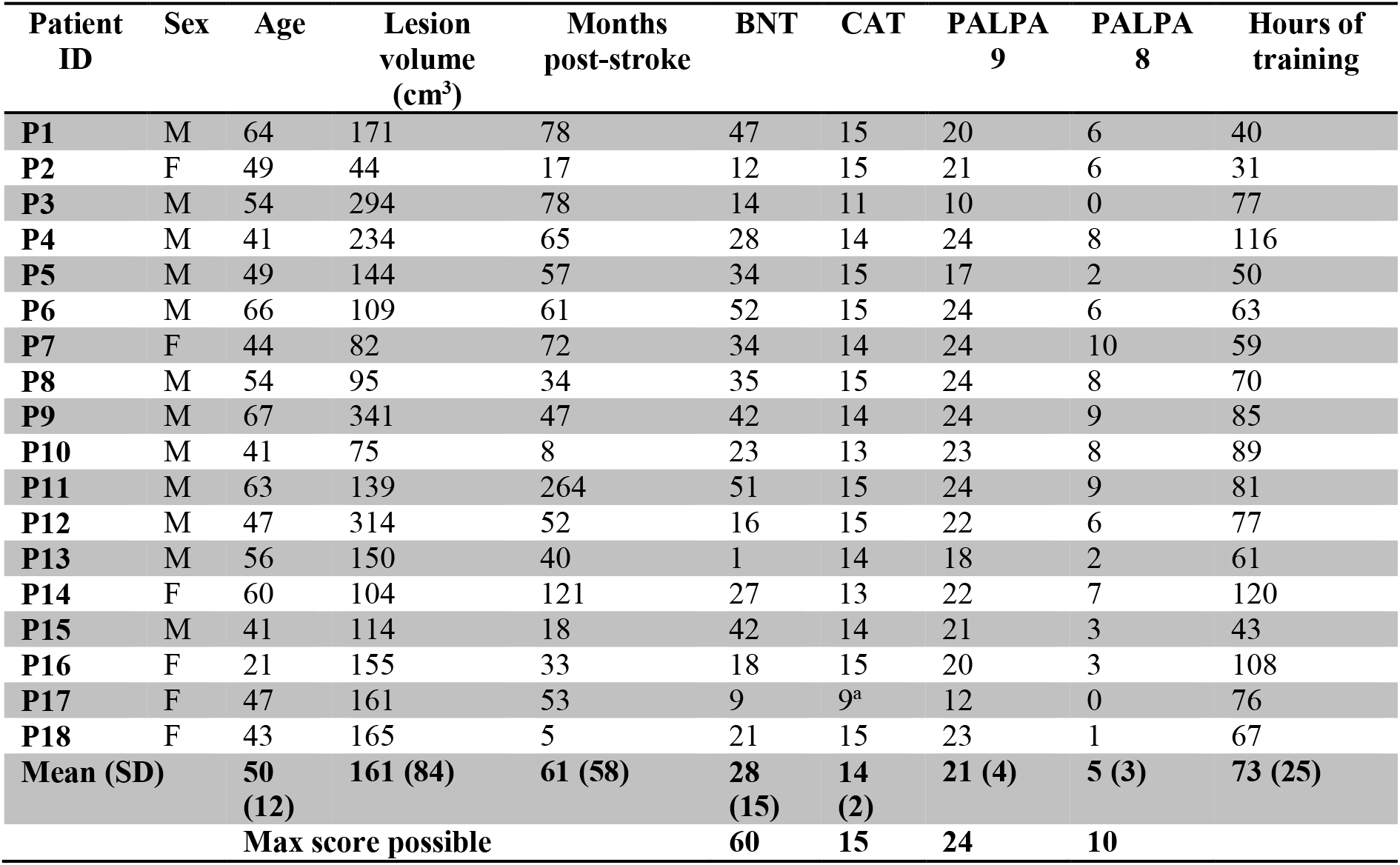
Demographic and clinical data of the patients

TRIPOD checklist for prediction model development.

| Section/Topic |  | Checklist Item | Page |
| --- | --- | --- | --- |
| **Title and abstract** |  |  |  |
| Title | 1 | Identify the study as developing and/or validating a multivariable prediction model, the target population, and the outcome to be predicted. | X |
| Abstract | 2 | Provide a summary of objectives, study design, setting, participants, sample size, predictors, outcome, statistical analysis, results, and conclusions. | X |
| **Introduction** |  |  |  |
| Background and objectives | 3a | Explain the medical context (including whether diagnostic or prognostic) and rationale for developing or validating the multivariable prediction model, including references to existing models. | x |
|  | 3b | Specify the objectives, including whether the study describes the development or validation of the model or both. | x |
| **Methods** |  |  |  |
| Source of data | 4a | Describe the study design or source of data (e.g., randomized trial, cohort, or registry data), separately for the development and validation data sets, if applicable. | x |
|  | 4b | Specify the key study dates, including start of accrual; end of accrual; and, if applicable, end of follow-up. | X |
| Participants | 5a | Specify key elements of the study setting (e.g., primary care, secondary care, general population) including number and location of centres. | x |
|  | 5b | Describe eligibility criteria for participants. | X |
|  | 5c | Give details of treatments received, if relevant. | X |
| Outcome | 6a | Clearly define the outcome that is predicted by the prediction model, including how and when assessed. | X |
|  | 6b | Report any actions to blind assessment of the outcome to be predicted. | X |
| Predictors | 7a | Clearly define all predictors used in developing or validating the multivariable prediction model, including how and when they were measured. | X |
|  | 7b | Report any actions to blind assessment of predictors for the outcome and other predictors. | X |
| Sample size | 8 | Explain how the study size was arrived at. | x |
| Missing data | 9 | Describe how missing data were handled (e.g., complete-case analysis, single imputation, multiple imputation) with details of any imputation method. | x |
| Statistical analysis methods | 10a | Describe how predictors were handled in the analyses. | X |
|  | 10b | Specify type of model, all model-building procedures (including any predictor selection), and method for internal validation. | X |
|  | 10d | Specify all measures used to assess model performance and, if relevant, to compare multiple models. | X |
| Risk groups | 11 | Provide details on how risk groups were created, if done. | X |
| **Results** |  |  |  |
| Participants | 13a | Describe the flow of participants through the study, including the number of participants with and without the outcome and, if applicable, a summary of the follow-up time. A diagram may be helpful. | X |
|  | 13b | Describe the characteristics of the participants (basic demographics, clinical features, available predictors), including the number of participants with missing data for predictors and outcome. | X |
| Model development | 14a | Specify the number of participants and outcome events in each analysis. | X |
|  | 14b | If done, report the unadjusted association between each candidate predictor and outcome. | X |
| Model specification | 15a | Present the full prediction model to allow predictions for individuals (i.e., all regression coefficients, and model intercept or baseline survival at a given time point). | X |
|  | 15b | Explain how to use the prediction model. | X |
| Model performance | 16 | Report performance measures (with CIs) for the prediction model. | X |
| **Discussion** |  |  |  |
| Limitations | 18 | Discuss any limitations of the study (such as nonrepresentative sample, few events per predictor, missing data). | X |
| Interpretation | 19b | Give an overall interpretation of the results, considering objectives, limitations, and results from similar studies, and other relevant evidence. | X |
| Implications | 20 | Discuss the potential clinical use of the model and implications for future research. | X |
| **Other information** |  |  |  |
| Supplementary information | 21 | Provide information about the availability of supplementary resources, such as study protocol, Web calculator, and data sets. | x |
| Funding | 22 | Give the source of funding and the role of the funders for the present study. | x |

